# JIB-04 has broad-spectrum antiviral activity and inhibits SARS-CoV-2 replication and coronavirus pathogenesis

**DOI:** 10.1101/2020.09.24.312165

**Authors:** Juhee Son, Shimeng Huang, Qiru Zeng, Traci L. Bricker, James Brett Case, Jinzhu Zhou, Ruochen Zang, Zhuoming Liu, Xinjian Chang, Houda H. Harastani, Lu Chen, Maria Florencia Gomez Castro, Yongxiang Zhao, Hinissan P. Kohio, Gaopeng Hou, Baochao Fan, Beibei Niu, Rongli Guo, Paul W. Rothlauf, Adam L. Bailey, Xin Wang, Pei-Yong Shi, Elisabeth D. Martinez, Sean P.J. Whelan, Michael S. Diamond, Adrianus C.M. Boon, Bin Li, Siyuan Ding

**Author notes:** These authors contributed equally: Juhee Son, Shimeng Huang, Qiru Zeng. Corresponding authors: Bin Li, Siyuan Ding.

## Abstract

Pathogenic coronaviruses represent a major threat to global public health. Here, using a recombinant reporter virus-based compound screening approach, we identified several small-molecule inhibitors that potently block the replication of the newly emerged severe acute respiratory syndrome virus 2 (SARS-CoV-2). Among them, JIB-04 inhibited SARS-CoV-2 replication in Vero E6 cells with an EC_50_ of 695 nM, with a specificity index of greater than 1,000. JIB-04 showed *in vitro* antiviral activity in multiple cell types against several DNA and RNA viruses, including porcine coronavirus transmissible gastroenteritis virus. In an *in vivo* porcine model of coronavirus infection, administration of JIB-04 reduced virus infection and associated tissue pathology, which resulted in improved weight gain and survival. These results highlight the potential utility of JIB-04 as an antiviral agent against SARS-CoV-2 and other viral pathogens.

## INTRODUCTION

The coronavirus disease 2019 (COVID-19) pandemic has caused unprecedented global morbidity, mortality, and socioeconomic destabilization. Thus, there is an urgent unmet need to develop safe and effective countermeasures to combat the disease beyond vaccine protection and provide immediate treatment. Multiple efforts are underway to identify candidate drugs that inhibit the replication of severe acute respiratory syndrome virus 2 (SARS-CoV-2) (Riva *et al*., 2020; Touret *et al*., 2020; Dittmar *et al*., 2021; Heiser *et al*., 2020; Mirabelli *et al*., 2020), the cause of COVID-19 (Wu *et al*., 2020; Zhou *et al*.,2020). So far, several small-molecule inhibitors that interfere with SARS-CoV-2 cell entry have been identified, including transmembrane serine protease inhibitors camostat (Hoffmann *et al*., 2020) and nafamostat (Wang *et al*., 2020a), and endosomal inhibitors including chloroquine and its derivatives (Wang *et al*., 2020a), E-64d (Hoffmann *et al*.,2020), apilimod (Kang *et al*., 2020), and 25-hydroxycholesterol (Zang *et al*., 2020a). Drug screens and structural studies also revealed compounds that target the viral enzymes of SARS-CoV-2, namely the RNA-dependent RNA polymerase (Yin *et al*., 2020; Gao *et al*., 2020; Kirchdoerfer and Ward, 2019; Nguyen *et al*., 2020; Sheahan *et al*., 2020) and the main protease (M^pro^, also known as 3CL^pro^) (Zhang *et al*., 2020; Dai *et al*., 2020; Jin *et al*.,2020; Nguyen *et al*., 2020). Here, utilizing a fluorescent SARS-CoV-2 virus and an imaging-based screen approach, we identified several known and previously unknown antiviral compounds that inhibit SARS-CoV-2 replication.

## RESULTS

To identify small molecules with anti-SARS-CoV-2 activity, we performed a screen using a recombinant SARS-CoV-2 that encoded mNeonGreen as a reporter of infection (Xie *et al*., 2020) and an in-house collection of ~200 compounds that comprised FDA-approved drugs, well-defined broad-spectrum antiviral agents, and investigational new drugs. We identified 157 compounds that had greater antiviral efficacy (>44.8% inhibition) than either chloroquine or remdesivir against SARS-CoV-2 replication in Vero E6 cells (**Fig. 1A** and **Dataset S1**). One of these drugs was a pan-Jumonji histone demethylase inhibitor 5-chloro-N-[(E)-[phenyl(pyridin-2-yl)methylidene]amino]pyridin-2-amine (JIB-04 E-isomer) (Wang *et al*., 2013) (**Fig. S1A**). We selected JIB-04 (JIB-04 E-isomer, unless noted otherwise) for further characterization because several histone demethylases were recently discovered as SARS-CoV-2 host dependency factors (Wei *et al*., 2021; Wang *et al*., 2021; Schneider *et al*., 2021) and JIB-04 has not been reported as an antiviral molecule, despite its established anti-tumor activity (Wang *et al*., 2013; Kim *et al*., 2018; Parrish *et al*., 2018; Bayo *et al*., 2018; Dalvi *et al*., 2017).

**Fig. 1.**
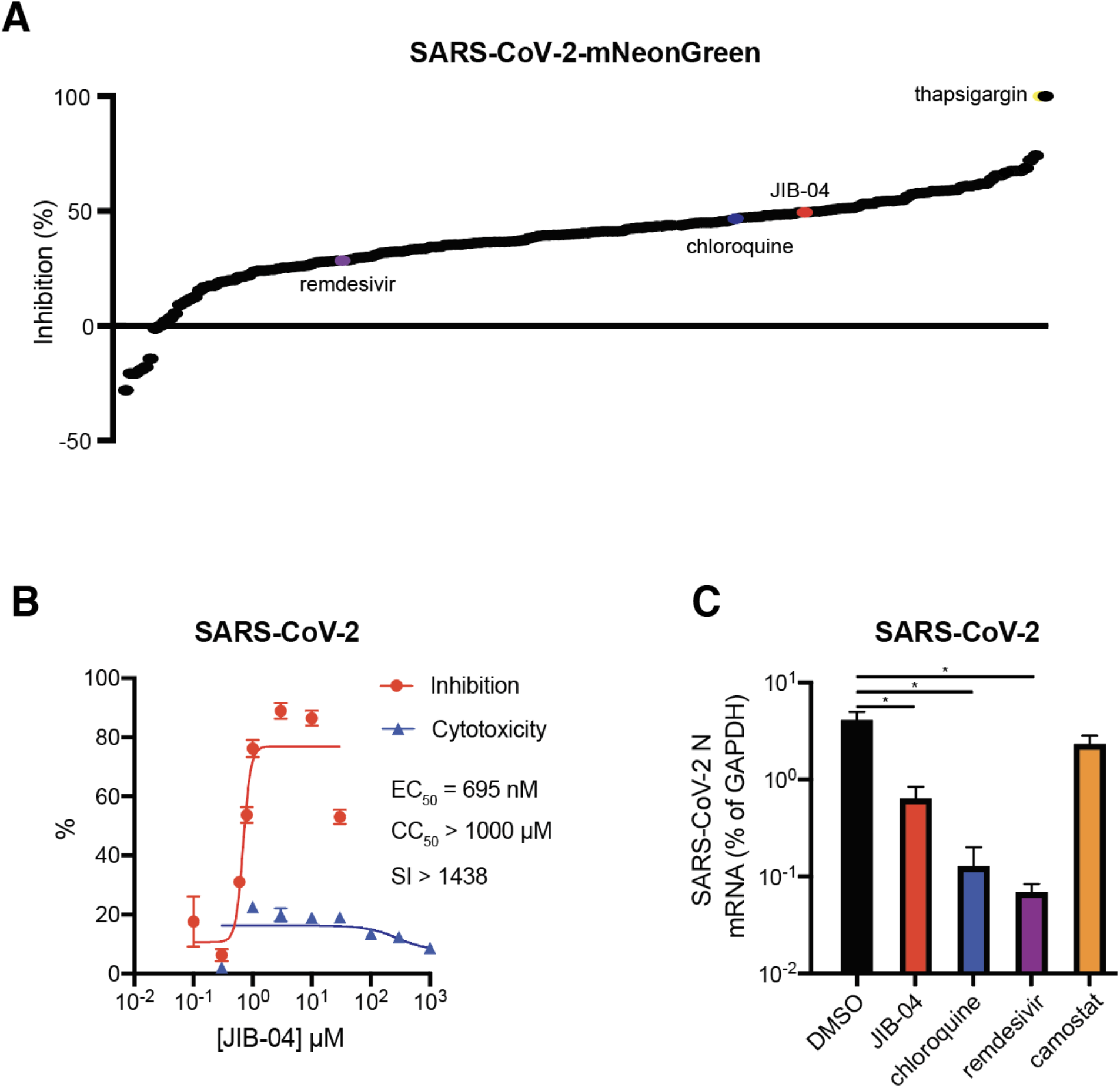
JIB-04 inhibits SARS-CoV-2 replication. (A) Small molecule inhibitor screen. Vero E6 cells were treated with individual compounds (listed in Table S1) at 10 μM for 1 hour (h) and infected with SARS-CoV-2-mNeonGreen (MOI=0.5). At 24 h post infection (hpi), cells were fixed, and nuclei were stained by Hoechst 33342. The intensities of mNeonGreen and Hoechst were quantified by the Typhoon biomolecular imager and the Cytation plate reader, respectively. The ratio of mNeonGreen and Hoechst is plotted as percentage of inhibition. (B) Dose-response curve of wild-type SARS-CoV-2 replication with JIB-04 treatment. Vero E6 cells were treated with JIB-04 for 1 h and infected with a clinical isolate of SARS-CoV-2 (MOI=0.5). S protein levels were quantified at 24 hpi based on immunofluorescence. For CC_50_ measurement, cells were treated with JIB-04 at 0.3 μM to 1 mM for 25 h. SI: selectivity index. (C) Intracellular viral RNA levels of cells treated with compounds and subsequently infected with wild-type SARS-CoV-2. Vero E6 cells were treated with JIB-04 (10 μM), chloroquine (10 μM), remdesivir (3 μM), or camostat (10 μM) for 1 h and infected with a clinical isolate of SARS-CoV-2 (MOI=0.5). SARS-CoV-2 RNA levels at 24 hpi were measured by RT-qPCR. For all panels except A, experiments were repeated at least three times with similar results. Fig. 1A was performed once with raw data included in Dataset S1. Data are represented as mean ± SEM. Statistical significance is from pooled data of the multiple independent experiments (*p≤0.05).

We tested whether JIB-04 treatment could inhibit replication of a clinical isolate of SARS-CoV-2 (2019-nCoV/USA-WA1/2020 strain). Viral antigen staining showed that a 1-hour pre-treatment with JIB-04 suppressed SARS-CoV-2 infection in Vero E6 cells with an EC_50_ value of 695 nM (95% confidence interval of 567-822 nM) (**Fig. 1B**). Cell viability did not fall below 50% even at 1 mM of JIB-04 treatment, making the selectivity index of JIB-04 higher than 1,000. Intracellular SARS-CoV-2 RNA levels also were reduced significantly by JIB-04, but not by camostat, a TMPRSS serine protease inhibitor (**Fig. 1C**).

To examine whether JIB-04 targets SARS-CoV-2 spike protein-mediated entry or other post-entry pathways (*e.g*., translation, replication, or assembly) shared between SARS-CoV-2 and other viruses, we tested JIB-04 against vesicular stomatitis virus (VSV) that expresses eGFP as a marker of infection (Cherry *et al*., 2005) and a replication-competent chimeric VSV in which the native glycoprotein (G) was replaced by the spike of SARS-CoV-2 (VSV-eGFP-SARS-CoV-2) (Case *et al*., 2020). JIB-04 suppressed replication of both viruses in MA104 and Vero E6-TMPRSS2 cells (**Fig. 2A-B**). Flow cytometry analysis of cells at 6h post-infection revealed a reduction in eGFP expression demonstrating that the inhibitory effect of JIB-04 occurs during either entry or gene-expression (**Fig. 2B**). Virus-infected cells also showed less GFP intensity with JIB-04 treatment (**Fig. S2A**). JIB-04 inhibited VSV-SARS-CoV-2 infection dose-dependently without apparent cytotoxicity (**Fig. 2C** and **S2B**), which became more apparent when cells were inoculated with virus at a low multiplicity of infection (MOI) (**Fig. S2C**). At 30 μM, JIB-04 treatment resulted in a 100-fold reduction of intracellular VSV-SARS-CoV-2 RNA levels (**Fig. S2D**).

**Fig. 2.**
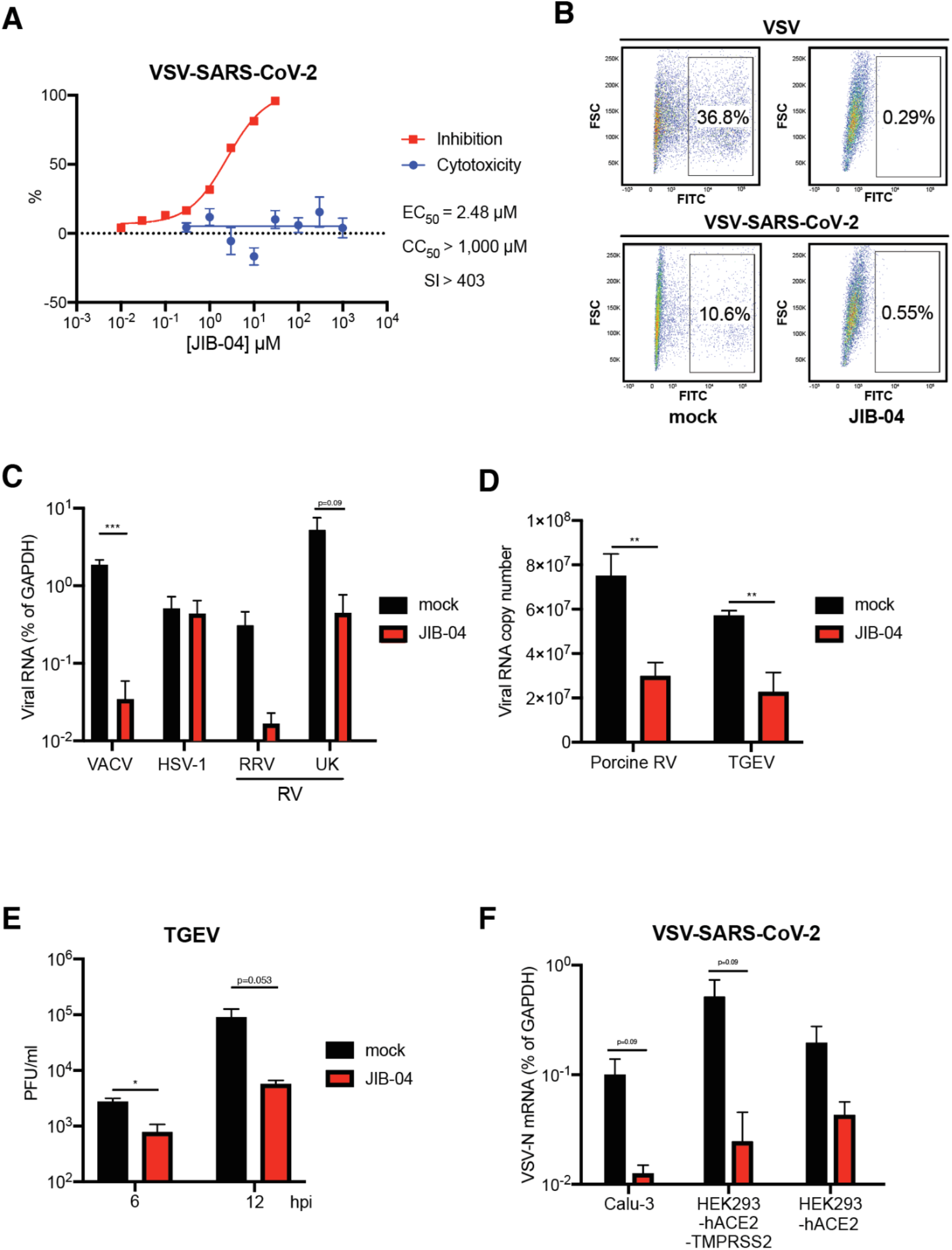
JIB-04 broadly inhibits DNA and RNA viruses in different cell types. (A) Dose-response analysis of VSV-SARS-CoV-2 replication and cytotoxicity with JIB-04 treatment. For EC_50_ measurement, MA104 cells were treated with compounds at 0.01 to 30 μM for 1 h and infected with VSV-SARS-CoV-2 (MOI=3) for 24 h. For CC_50_ measurement, cells were treated with compounds at 0.1 μM to 3 mM for 25 h. SI: selectivity index. (B) Virus infectivity with JIB-04 treatment. Vero E6-TMPRSS2 cells were treated with compounds (10 μM) for 1 h and infected with VSV or VSV-SARS-CoV-2 (MOI=3). At 6 hpi, percentages of GFP positive cells were quantified by flow cytometry. (C) Intracellular viral RNA levels with JIB-04 treatment. MA104 cells were treated with compounds (10 μM) for 1 h and infected with vaccinia virus (VACV), herpes simplex virus-1 (HSV-1), or rotavirus (RV, RRV and UK strains) (MOI=1). Viral RNA levels at 24 hpi were measured by RT-qPCR for VACV B10R, HSV-1 ICP-27, and RV NSP5, respectively. (D) Viral RNA copy numbers with JIB-04 treatment. HEK293 cells were treated with JIB-04 (10 μM) for 6 h and infected with porcine rotavirus (MOI=0.01) for 6 h. ST cells were treated with JIB-04 (10 μM) for 12 h and infected with transmissible gastroenteritis virus (TGEV) (MOI=0.01) for 12 h. Viral RNA copy numbers were measured by RT-qPCR. (E) TGEV titers in the cell supernatant with JIB-04 treatment. ST cells were treated with JIB-04 (10 μM) for 12 h and infected with TGEV (MOI=0.01). Virus titers at 6 and 12 hpi were measured by plaque assays. (F) Intracellular viral RNA levels with JIB-04 treatment in different cell types. Calu-3 cells, HEK293-hACE2 and HEK293-hACE2-TMPRSS2 were treated with compounds (10 μM) for 1 h and infected with VSV-SARS-CoV-2 (MOI=1). VSV RNA levels at 24 hpi were measured by RT-qPCR. All experiments were repeated at least three times with similar results. Data are represented as mean ± SEM. Statistical significance is from pooled data of the multiple independent experiments (*p≤0.05; **p≤0.01; ***p≤0.001).

We next evaluated the_antiviral activity of JIB-04 against other viruses. Though JIB-04 did not diminish replication of herpes simplex virus 1, it inhibited the replication of vaccinia virus, another DNA virus, and several strains of rotavirus (RV), a double-stranded RNA virus (RV) (**Fig. 2C-D**). JIB-04 also inhibited the replication of transmissible gastroenteritis virus (TGEV) (**Fig. 2D-E** and **Fig. S2E**), a porcine coronavirus that infects the small intestine of pigs and causes lethal diarrhea (Saif, 2004). This indicates that the antiviral effect of JIB-04 is not limited to single-stranded RNA viruses in cell culture.

Although JIB-04 inhibits the replication of both SARS-CoV-2 and VSV-SARS-CoV-2 in monkey kidney epithelial cell lines, a primary *in vivo* target of SARS-CoV-2 is ciliated airway epithelial cells (Hou *et al*., 2020). We therefore examined the inhibitory effect if JIB-04 on SARS-CoV-2 infection of the human lung epithelial cell line Calu-3 (Hoffmann *et al*., 2020; Sheahan *et al*., 2020). We validated that JIB-04 retained its antiviral activity against VSV-SARS-CoV-2 in Calu-3 cells (**Fig. 2G**). VSV-SARS-CoV-2 replication was also inhibited by JIB-04 in HEK293 cells ectopically expressing human ACE2, an entry receptor for SARS-CoV-2 (Hoffmann *et al*., 2020), with or without ectopic TMPRSS2 expression (**Fig. 2G**).

We next sought to understand the mechanisms of antiviral activity of JIB-04. Although JIB-04 has been previously connected to interferon (IFN) and autophagy activation (Wang *et al*., 2013; Xu *et al*., 2018), the antiviral activity that we observed was independent of these pathways. JIB-04 treatment did not lead to the induction of IFN and IFN-stimulated gene expression or the formation of LC3-positive punctate structures (**Fig. S3A-B**). To explore the mechanisms of antiviral action, we utilized a drug combination approach. When a combination of JIB-04 with chloroquine was evaluated based on the highest single agent (HAS) synergy model with SynergyFinder 2.0 (Ianevski *et al*., 2020), JIB-04 was shown to exert a synergistic antiviral effect with chloroquine in MA104 cells (**Fig. 3A**). A combination of JIB-04 and camostat also was synergistically antiviral in Calu-3 cells (**Fig. S3C**), indicating that JIB-04 likely targets a different pathway.

**Fig. 3.**
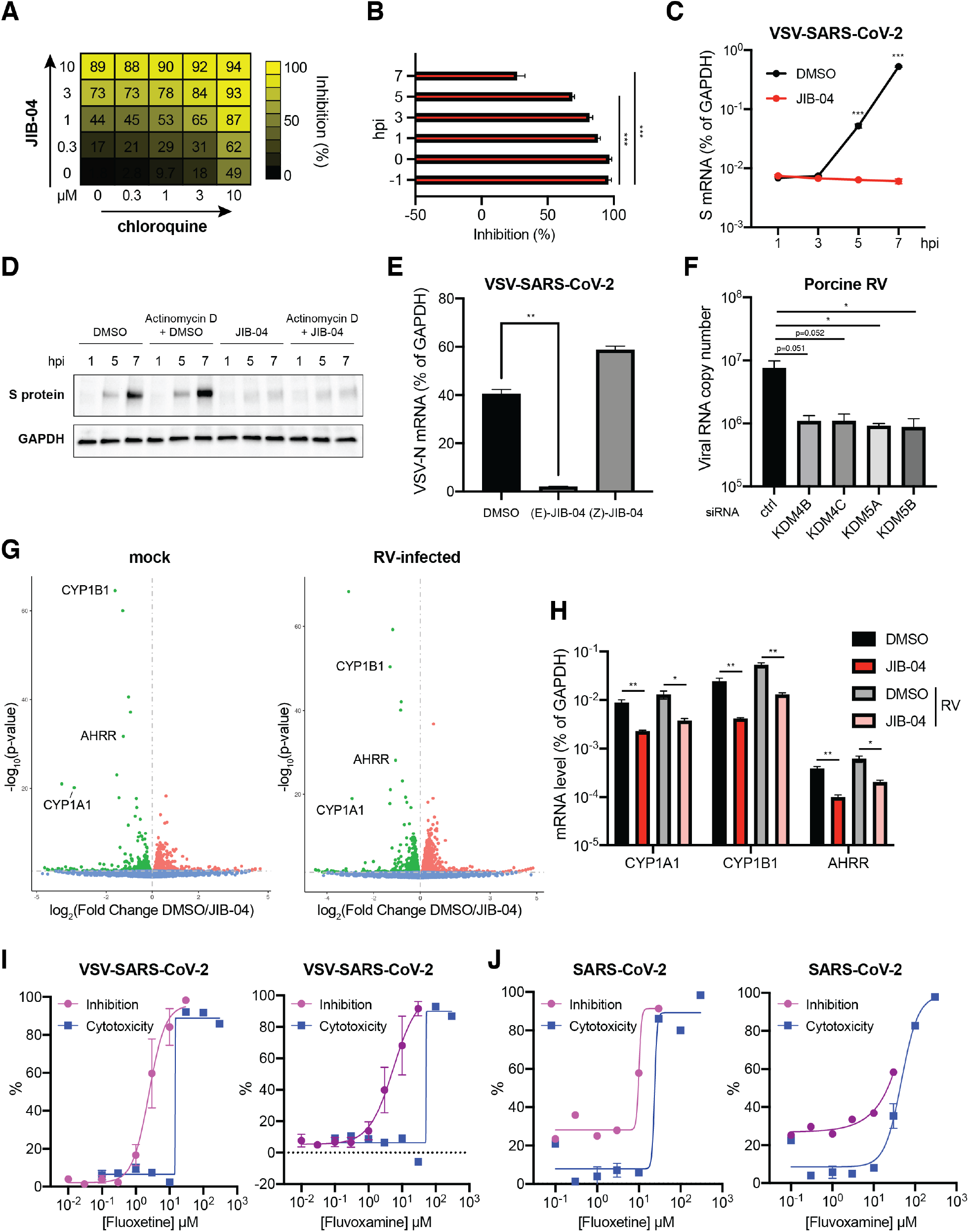
JIB-04 exhibits distinct post-entry antiviral mechanisms. (A) Drug combination dose-response matrix and VSV-SARS-CoV-2 replication. MA104 cells were treated with JIB-04 and chloroquine for 1 h and infected with VSV-SARS-CoV-2 (MOI=3). GFP signals at 24 hpi were quantified to calculate the percentage of inhibition. (B) Time of compound addition and VSV-SARS-CoV-2 replication. MA104 cells were treated with JIB-04 (10 μM) at indicated time points relative to VSV-SARS-CoV-2 infection (MOI=3, 0 hpi). GFP signals at 8 hpi were quantified to calculate the percentage of inhibition. (C) Intracellular SARS-CoV-2 S RNA levels with JIB-04 treatment. MA104 cells were treated with JIB-04 (10 μM) for 1 h and infected with VSV-SARS-CoV-2 (MOI=1) for 1, 3, 5, and 7 h. S RNA levels were measured by RT-qPCR. (D) Western blot analysis of SARS-CoV-2 S protein levels with JIB-04 treatment. MA104 cells were treated with JIB-04 (10 μM) for 1 h and infected with VSV-SARS-CoV-2 (MOI=1) for 1, 5, and 7 h. For Actinomycin D, 10 μg/ml actinomycin D was added to the media 15 min before DMSO or JIB-04 treatment. (E) Intracellular viral RNA levels of cells treated with JIB-04 E-isomer or Z-isomer and subsequently infected with VSV-SARS-CoV-2. MA104 cells were treated with JIB-04 isomer (10 μM) for 1 h and infected with VSV-SARS-CoV-2 (MOI=1). VSV-N levels at 24 hpi were measured by RT-qPCR. (F) Histone demethylase siRNA knockdown and RV replication. HEK293 cells were transfected with scrambled siRNA or siRNA targeting indicated histone demethylases for 48 h and infected with porcine RV (MOI=0.01). Viral RNA copy numbers at 12 hpi were quantified by RT-qPCR. (G) Volcano plot of differentially expressed transcripts with JIB-04 treatment and RV infection. HEK293 cells were treated with DMSO or JIB-04 (10 μM) for 12 h, and mock-infected (left panel) or infected with porcine RV (MOI=0.01, right panel) for another 12 h. Red dots represent upregulated genes and green dots represent downregulated genes in JIB-04 treated cells. (H) Expression of three top genes in (G) with JIB-04 treatment. HEK293 cells were treated with JIB-04 (10 μM) for 12 h and mock-infected or infected porcine RV (MOI=0.01) for 12 h. mRNA levels of *CYP1A1, CYP1B1, and AHRR* at 12 hpi were measured by RT-qPCR. (I) Dose-response analysis of VSV-SARS-CoV-2 replication with fluoxetine or fluvoxamine treatment. MA104 cells were treated with compounds at 0.01 to 30 μM for 1 h and infected with VSV-SARS-CoV-2 (MOI=3). GFP signals at 24 hpi were quantified to calculate the percentage of inhibition. For CC_50_ measurement, cells were treated with compounds at 0.1 μM to 300 μM for 25 h. (J) Dose-response analysis of wild-type SARS-CoV-2 replication with fluoxetine or fluvoxamine treatment. Vero E6 cells were treated with compounds for 1 h and infected with a clinical isolate of SARS-CoV-2 (MOI=0.5). S protein levels at 24 hpi were quantified based on immunofluorescence. For CC_50_ measurement, cells were treated with compounds at 0.1 μM to 300 μM for 25 h. For all panels except A and J, experiments were repeated at least three times with similar results. Fig. 3A was performed twice. Inhibition assay in Fig. 3J was performed once and cytotoxicity assay was performed in triplicates. Data are represented as mean ± SEM. Statistical significance is from pooled data of the multiple independent experiments (*p≤0.05; **p≤0.01; ***p≤0.001).

A possible antiviral role for JIB-04 at a post-entry step was supported by the time of addition experiments (**Fig. 3B**). A 1-h pre-treatment of cells with JIB-04 reduced SARS-CoV-2 spike mRNA transcription following VSV-SARS-CoV-2 infection (**Fig. 3C**), and translation of newly synthesized spike protein, which could not be achieved with Actinomycin D treatment (**Fig. 3D**). These results suggest that JIB-04 might repress virus replication by interfering with the viral RNA transcription or stability.

We also assessed whether the antiviral activity of JIB-04 is linked to its epigenetic modulatory action. Unlike its E-isomer, the Z-isomer of JIB-04 does not inhibit histone demethylases at similar doses (Wang *et al*., 2013). When we compared the antiviral efficacy of these two JIB-04 isomers against VSV-SARS-CoV-2 in MA104 cells, the Z-isomer did not inhibit the replication of virus (**Fig. 3E)**. The disparity between the isomers suggests that epigenetic enzyme inhibition is involved in antiviral mechanisms of JIB-04. To examine the cellular pathways modulated by JIB-04, we performed small interfering RNA (siRNA)-mediated knockdown of JIB-04 cellular targets (*i.e*., histone demethylases KDM4B, KDM4C, KDM5A, or KDM5B (Wang *et al*., 2013)). Knockdown of each gene successfully recapitulated the antiviral effect of JIB-04 (**Fig. 3F** and **Fig. S3E**). These results led us to hypothesize that JIB-04 treatment promoted H3K9 and H3K27 methylation and silenced expression of a subset of genes, triggering the antiviral effect. To identify potential target genes, we performed RNA-sequencing on the cells pre-treated with vehicle or JIB-04 with or without virus infection (**Fig. 3G**). Pathway analysis revealed dampened metabolic signaling pathways such as cytochrome P450 system in JIB-04 treated cells (**Fig. S3F**). Specifically, JIB-04 treatment downregulated two cytochrome P450 enzymes, *CYP1A1* and *CYP1B1*, and aryl hydrocarbon receptor repressor (*AHRR*),which represses transactivator of *CYP1A1* and *CYP1B1* (Karchner *et al*., 2002). We validated by quantitative PCR that JIB-04 treatment reduced *CYP1A1, CYP1B1*, and *AHRR* mRNA levels by 4-6-fold (**Fig. 3H**). To explore the pharmacological utility of this finding, we tested the antiviral activity of cytochrome P450 enzyme inhibitors fluoxetine and fluvoxamine (Hemeryck and Belpaire, 2002). Both compounds inhibited the replication of VSV-SARS-CoV-2 (**Fig. 3I**) and wild-type SARS-CoV-2 (**Fig. 3J**).

Given that JIB-04 prevents coronavirus replication *in vitro*, we used a neonatal pig TGEV infection model (Luo *et al*., 2019) to assess the efficacy of JIB-04 against coronavirus infection *in vivo*. Two-day old piglets were injected via an intraperitoneal route with JIB-04 twice before the oral inoculation of TGEV (**Fig. 4A**). We monitored body weight on a daily basis and recorded diarrhea development and mortality every 6 h. The animals in the control group lost more weight and had more severe diarrhea than those receiving JIB-04 (**Fig. S4A-B**). At 2 days post infection, 3 of 5 piglets in the DMSO group succumbed to infection as compared to 1 out of 5 animals in the JIB-04 group (**Fig. 4B**). Consistent with our *in vitro* results (**Fig. 2E-F**), the TGEV viral burden throughout the gastrointestinal tract was substantially lower in the JIB-04 treated group (**Fig. 4C-D**). JIB-treated animals also had lower number of viral antigen positive cells in their intestinal epithelium (**Fig. S4C**) and showed less enteropathy than the control group (**Fig. 4E**). Taken together, our data demonstrate *in vivo* antiviral activity of JIB-04 against a porcine coronavirus.

**Fig. 4.**
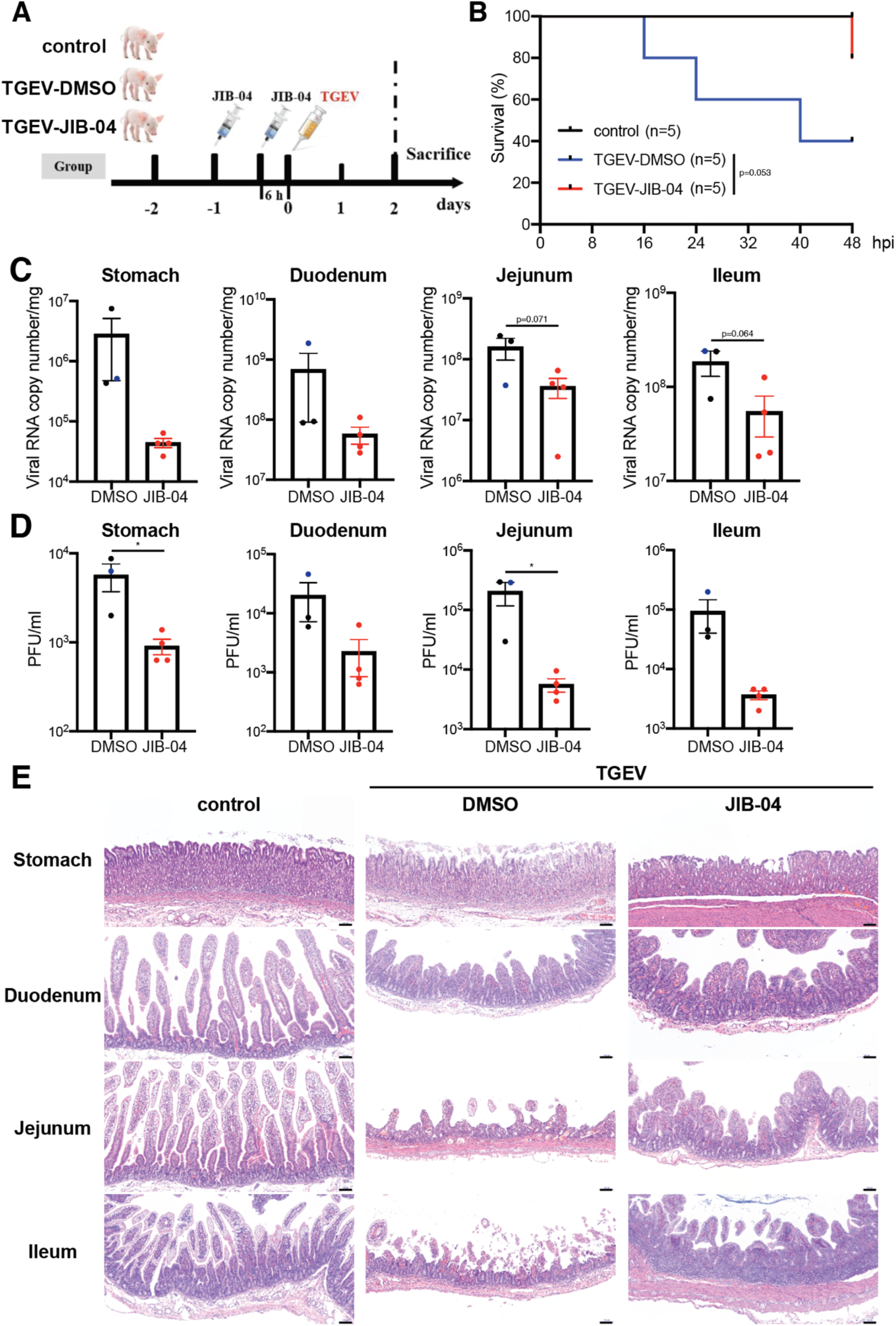
JIB-04 suppresses TGEV replication and pathogenesis in pigs. (A) Experimental schemes for testing the protective efficacy of JIB-04 treatment against TGEV challenge in three groups of neonatal pigs. Control: DMSO injection, mock infection; TGEV-DMSO: DMSO injection, TGEV infection; TGEV-JIB-04: JIB-04 injection, TGEV infection. (B) Survival curve of TGEV infected pigs with JIB-04 treatment. Neonatal pigs were intraperitoneally injected with vehicle control DMSO or JIB-04 and mock-infected or infected with 1.2 × 10^7^ PFU of TGEV. Survival was monitored every 8 h with data censored at 48 hpi, when all pigs were sacrificed. (C) TGEV RNA copy numbers in the gastrointestinal (GI) tract of infected pigs. TGEV infected piglets were sacrificed at 48 hpi. For the DMSO group, two animals sacrificed at 48 hpi and one that died at 40 hpi (colored in blue) were examined. For the JIB-04 groups, four animals sacrificed at 48 hpi were examined. TGEV genome copy numbers at 48 hpi were quantified by RT-qPCR. (D) Same as (C) except that virus titers were measured by plaque assays. (E) Hematoxylin and eosin staining of different GI tract sections from pigs sacrificed at 48 hpi. Representative images of 3 animals. Scale bar, 100 μm. Data are represented as mean ± SEM. Statistical significance is from pooled data of the multiple independent experiments (*p≤0.05).

## DISCUSSION

Using a repurposed compound screening approach, we identified drugs with reported inhibitory activity against SARS-CoV-2, such as tetrandrine (Ou *et al*., 2020) and arbidol (Wang *et al*., 2020b). We also characterized a number of small-molecules (JIB-04, AG-1478, nigericin, etc.) without known antiviral activity as inhibitors of SARS-CoV-2 infection. While this manuscript was in preparation, a new study identified thapsigargin, the compound that showed the highest anti-SARS-CoV-2 activity in our screen as a broad antiviral against coronavirus (Al-Beltagi *et al*., 2021), which validates our screen approach. Notably, several top hit compounds in the screen converge on the endosomal trafficking pathway: brefeldin A, concanamycin A, tetrandrine, and U18666A. Furthermore, FTY720 induced formation of enlarged endosome/lysosome structure, similar to that triggered by apilimod treatment (Kang *et al*., 2020). All of these point to an important role of endosomal trafficking in SARS-CoV-2 entry and infection, at least in cell culture.

Our results highlight JIB-04 as a potential therapeutic for SARS-CoV-2 and suggest further evaluation of this drug that has mainly been associated with its anti-cancer activities. Another compound in our screen, GSK-J4, also is a histone demethylase inhibitor that targets KDM6B. However, unlike JIB-04, GSK-J4 failed to reduce viral burden in Vero E6 cells upon SARS-CoV-2 infection to an extent comparable to chloroquine. Thus, we speculate that there might be specific roles played by certain KDM family members in the interactions between the host and SARS-CoV-2.

### Limitations of the study

We have tried to be careful about establishing the broad-spectrum antiviral activity of JIB-04. Indeed, we have shown examples from distinct virus families (double-strand DNA virus: vaccinia virus; single-strand positive-strand RNA virus: SARS-CoV-2 and TGEV; single-strand negative-strand RNA virus: VSV and VSV-SARS-CoV-2; double-strand RNA virus: rotavirus). However, we have yet to test single-strand DNA viruses and retroviruses. We also did not examine whether JIB-04 has antiviral activity against the newly emerging SARS-CoV-2 variants. We showed that JIB-04 modulates cytochrome P450 genes and targeting these genes by well-established selective serotonin uptake inhibitor also led to the inhibition of SARS-CoV-2 replication. Nevertheless, we do not know how modulation of cytochrome P450 genes correlates with transcriptional repression of SARS-CoV-2 RNA that we observed after JIB-04 treatment. It is plausible that JIB-04 induces these two effects separately, which needs to be characterized in future studies. Lastly, though we provided evidence that JIB-04 protects against coronavirus infection *in vivo* using a porcine TGEV model, TGEV is an animal coronavirus that targets the enteric rather than the respiratory system. Protection against SARS-CoV-2 *in vivo* should be tested in a transgenic mouse or hamster model. Finally, while the distinct efficacy of the E vs Z isomers points to inhibition of Jumonji demethylases as contributing to the antiviral effects, direct evidence of this mechanism in multiple models would strengthen this conclusion.

## MATERIALS AND METHODS

### Reagents, cells, and viruses

#### Reagents

JIB-04 E-isomer used in *in vitro* assays (S7281, Selleckchem, 99.8% purity), JIB-04 E-isomer used in *in vivo* experiments (HY-13953, Med-ChemExpress), Actinomycin D (A5156, Sigma), Fluvoxamine maleate (S1336, Selleckchem), Fluoxetine HCl (S1333, Selleckchem), low molecular weight poly(I:C) complexed with LyoVec (tlrl-picwlv, InvivoGen). EGFP-LC3 plasmid was a gift from Christina Stallings at Washington University School of Medicine. pUC19 empty plasmid was used as mock in all transfection experiments. JIB-04 Z-isomer used in control experiments by synthesized as originally described (Wang et al, 2013).

#### Cells

Vero E6 cells (CRL-1586, ATCC) and Vero cells (CCL81, ATCC) were cultured in DMEM supplemented with 10% fetal bovine serum (FBS), 10 mM HEPES, 1 mM sodium pyruvate, 0.1 mM non-essential amino acids, and 1X Penicillin-Streptomycin-Glutamine. Calu-3 cells (HTB-55, ATCC) and swine ST cells (CRL-1746, ATCC) were DMEM supplemented with 10% FBS and 1 X Penicillin-Streptomycin-Glutamine. HEK293, HEK293-hACE2, and HEK293-hACE2-TMPRSS2 cells were cultured in complete DMEM containing G418 and/or blasticidin and used as previously described (Zang *et al*., 2020a). MA104 and Vero E6-TMPRSS2 cells were cultured as before (Zang *et al*., 2020b).

#### Viruses

Rhesus RV RRV strain, bovine RV UK strain, and porcine RV NJ2012 strain (Genbank: MT874983-MT874993) were propagated and titrated as before (Ding *et al*.,2018). Vaccinia virus MVA strain was used as before (Li *et al*., 2017). HSV-1 syn17+ strain was a gift from Ann Arvin at Stanford University. TGEV JS2012 strain was propagated as before (Guo *et al*., 2020). TGEV was titrated by serial dilutions in cells in 96-well plates that were seeded at a density of 1 × 10^4^ cells per well. Cytopathic effects were observed at 3-7 dpi and the TCID50 values were calculated and converted to PFU/ml. A clinical isolate of SARS-CoV-2 (2019-nCoV/USA-WA1/2020 strain) was obtained from the Centers for Disease Control and Prevention. A SARS-CoV-2 mNeonGreen reporter virus was used as previously reported (Xie *et al*., 2020). Both the clinical isolate and the mNeonGreen SARS-CoV-2 viruses were propagated in Vero CCL81 cells and titrated by focus-forming assays on Vero E6 cells. Recombinant VSV-eGFP (Cherry *et al*., 2005) and VSV-eGFP-SARS-CoV-2 were previously described (Case *et al*., 2020). Cells infected with viruses expressing GFP were imaged with an ECHO REVOLVE 4 fluorescence microscope. Plaque assays were performed in MA104 cells seeded in 6-well plates using an adapted version of the rotavirus plaque assay protocol (Ding *et al*., 2018).

### Inhibitor screen

The small-molecule inhibitors used in this study are from in-house collection and the COVID Box (Medicines for Malaria Venture; www.mmv.org/mmv-open/covid-box). Compound names, vendors, and catalog numbers are listed in **Table S1**. At 24 hpi, cells were fixed in 4% paraformaldehyde (PFA) in PBS and stained with Hoechst 33342. The levels of viral antigens, reflected by mNeonGreen signals, were scanned by Amersham Typhoon 5 (GE). Image background was subtracted using rolling ball algorithm (radius = 5 pixels). To minimize imaging artifacts and well-to-well variation, we removed the region which fell below the threshold calculated by Moments algorithm. The number of positive pixels and total intensity (after background subtraction) were computed for each well and log10 transformed. The number of cells in each well was quantified based on Hoechst 33342 staining detected by Cytation 3 imaging reader (BioTek). Image analysis was performed using ImageJ and customized R scripts. The quantification of mNeonGreen and Hoechst 33342 is provided in **Dataset S1**.

### Cell cytotoxicity assay

The viability of Vero E6 and MA104 cells after drug treatment was determined using the Cell Counting Kit 8 (ab228554, Abcam). Briefly, cells in 96-well plates were treated with JIB-04 at desired concentrations at 37°C. After a 25-h incubation, the inhibitor-containing medium was replaced with fresh complete medium with 10 μl of WST-8 solution in each well. The cells were incubated at 37 °C for 2 h with protection from light. Absorbance at 460nm was measured using Gen5 software and a BioTek ELx800 Microplate Reader.

### RNA extraction and quantitative PCR

Total RNA was extracted from cells using RNeasy Mini kit (Qiagen). For spike plasmid transfection experiments, total RNA was extracted using Aurum Total RNA Mini Kit (Biorad) with DNase digestion. Complementary DNA was synthesized with High Capacity cDNA Reverse Transcription kit (Thermo Fisher) as previously described (Bolen *et al*.,2013). Quantitative PCR was performed using AriaMX (Agilent) with 12.5 μl of either Power SYBR Green master mix or Taqman master mix (Applied Biosystems) in a 25 μl reaction. Gene expression was normalized to the housekeeping gene GAPDH. All SYBR Green primers and Taqman probes used in this study are listed in **Table S2**.

### Western blotting

Cells were lysed in RIPA buffer supplemented with protease inhibitor cocktail and phosphatase inhibitor. Lysates were boiled for 5 min in 1 x Laemmli Sample Buffer (Bio-Rad) containing 5% β-mercaptoethanol. Proteins were resolved in SDS-PAGE and detected as described(Ding *et al*., 2014) using the following antibodies: GAPDH (631402, Biolegend), rotavirus VP6 (rabbit polyclonal, ABclonal technology), and SARS-CoV-2 S2 (40592-T62, Sino Biological). Secondary antibodies were anti-rabbit (7074, Cell Signaling) or anti-mouse (7076, Cell Signaling) immunoglobulin G horseradish peroxidase-linked antibodies. Protein bands were visualized with Clarity ECL Substrate (Bio-rad) and a Biorad Gel Doc XR system.

### Small interfering RNA transfection

HEK293 cells were transfected using Lipofectamine RNAiMAX Transfection Reagent (Thermo Fisher). Cells were harvested at 48 h post transfection, and knockdown efficiency was determined by RT-qPCR. siRNA transfected cells were infected with PoRV (MOI=0.01) for 12 h and viral RNA copy numbers were examined by RT-qPCR. All siRNA used in this study were designed and synthesized by GenePharma (Shanghai, China) and their sequences of siRNAs are listed in **Table S2**.

### Flow cytometry

Vero E6-TMPRSS2 cells were inoculated with VSV-GFP or VSV-SARS-CoV-2 at an MOI of 3 for 1 h at 37°C. At 6 hpi, cells were harvested and fixed in 4% PFA in PBS. Percentage of GFP positive cells and GFP intensity were determined by BD LSRFortessa™ X-20 cell analyzer and analyzed by FlowJo v10.6.2 (BD).

### RNA-sequencing

HEK293 cells were pre-treated with JIB-04 (10 μM) for 12 h, and mock or porcine RV-infected (MOI=0.01) for another 12 h. Total RNA from cells in triplicate was extracted using RNeasy Mini kit (Qiagen). RNA sample quality was measured by both NanoDrop spectrophotometer (Thermo Fisher) and Bioanalyzer 2100 (Agilent). Libraries were sequenced on the Illumina NovaSeq 6000 platform. Differential gene expression analysis was performed using DESeq2. The RNA-seq raw and processed datasets were deposited onto NCBI Gene Expression Omnibus database (GSE156219).

### TGEV piglet infection

Newborn piglets (Landrace × Yorkshire) were spontaneously delivered from sows, and their body weights were recorded at birth. Fifteen neonatal male pigs at birth were obtained from a TGEV-free farm in Nanjing without suckling. All piglets were confirmed negative for TGEV by RT-PCR and ELISA (IDEXX, USA). The pigs were randomly separated into three groups, housed in separate rooms, and fed the same artificial milk substitutes that meet the nutrient and energy recommendations of the National Research Council [NRC, 2012] at the animal facility of the Institute of Veterinary Medicine, Jiangsu Academy of Agricultural Sciences, Nanjing, Jiangsu Province. The experiments were divided into three groups: a DMSO inoculation control group (control, n=5); a DMSO inoculation and TGEV infection group (TGEV-DMSO, n=5); a JIB-04 inoculation and TGEV infection group (TGEV-JIB-04, n=5)). Neonatal pigs were intraperitoneally injected twice with JIB-04 (75 mg/kg) or DMSO at 24 h and 6 h prior to TGEV infection. TGEV-DMSO and TGEV-JIB-04 groups were orally infected with 1×10^7.25^ (1.778×10^7^) TCID50 (equivalent to 1.245×10^7^ PFU) of TGEV in 1.5 ml of DMEM per pig. Neonatal pigs were weighed and observed for clinical signs every 8 h throughout the study. Serum samples were collected from each pig at 24 and 48 hpi to detect specific anti-TGEV antibodies. The occurrence of diarrhea was monitored, and its severity was recorded based on an established scoring system (Li *et al*., 2017). In brief, diarrhea was scored on the basis of color, consistency, and amount, and numbered as follows: 0 = normal; 1 = pasty; 2 = semi-liquid; 3 = liquid, and score ≥ 2 considered as diarrhea. At 48 hpi, all pigs were euthanized, and intestinal tissues were collected for pathological examination and viral load analysis using RT-qPCR and primers in **Table S2**.

### Histopathological and immunofluorescence analysis

Intestinal tissues harvested from pigs were fixed in 4% PFA in PBS and incubated in 50% ethanol overnight. After fixation, tissues were embedded in paraffin, sectioned, and subjected to hematoxylin and eosin staining by standard procedures. For immunofluorescence analysis, samples were incubated with rabbit anti-TGEV-N antibody (1:500, DA0224, Shanghai YouLong Biotech) for 30 min at 37 °C. After three washes, samples were stained with Cy3-conjugated goat anti-rabbit secondary antibody (Beyotime) and DAPI (Invitrogen). Images were obtained using a fluorescence microscope (Carl Zeiss).

### Ethics statement

Animal experiments were approved by the Committee on the Ethics of Animal Care and Use of the Science and Technology Agency of Jiangsu Province. The approval ID is NKYVET 2014-63, granted by the Jiangsu Academy of Agricultural Sciences Experimental Animal Ethics Committee. All efforts were made to minimize animal suffering. The virus challenge and tissue collection were performed in strict accordance with the guidelines of Jiangsu Province Animal Regulations (Decree No. 2020-18).

### Statistical analysis

All bar graphs were displayed as means ± standard error of mean (SEM). Statistical significance in data Fig. 2E, 2F, 3C, 4C, and 4D was calculated by Student’s *t* test using Prism 8.4.3 (GraphPad). Statistical significance in data Fig. 1C, 2D, 2G, 3B, 3E, 3G, S3A, S3C, and S4B was calculated by pairwise ANOVA using Prism 8.4.3. Non-linear regression (curve fit) was performed to calculate EC_50_ and CC_50_ values for Fig. 1B, 2A, and 2C using Prism 8.4.3. HSA synergy model was used to calculate the synergy scores of dose-response data in Fig. 3A. Gehan-Breslow-Wilcoxon test was used to compare the survival curves in Fig. 4B. All data were presented as asterisks (*p≤0.05; **p≤0.01; ***p≤0.001). All experiments other than Fig. 1A, 3I, and 4 were repeated at least twice. The raw data are included in **Table S3**.

## SUPPLEMENTARY MATERIALS

Table S1. List of chemicals used in the anti-SARS-CoV-2 compound screen

Table S2. List of qPCR primers and siRNA

Table S3. Raw data

## Supplemental Figures and Legends

**Fig. S1.**
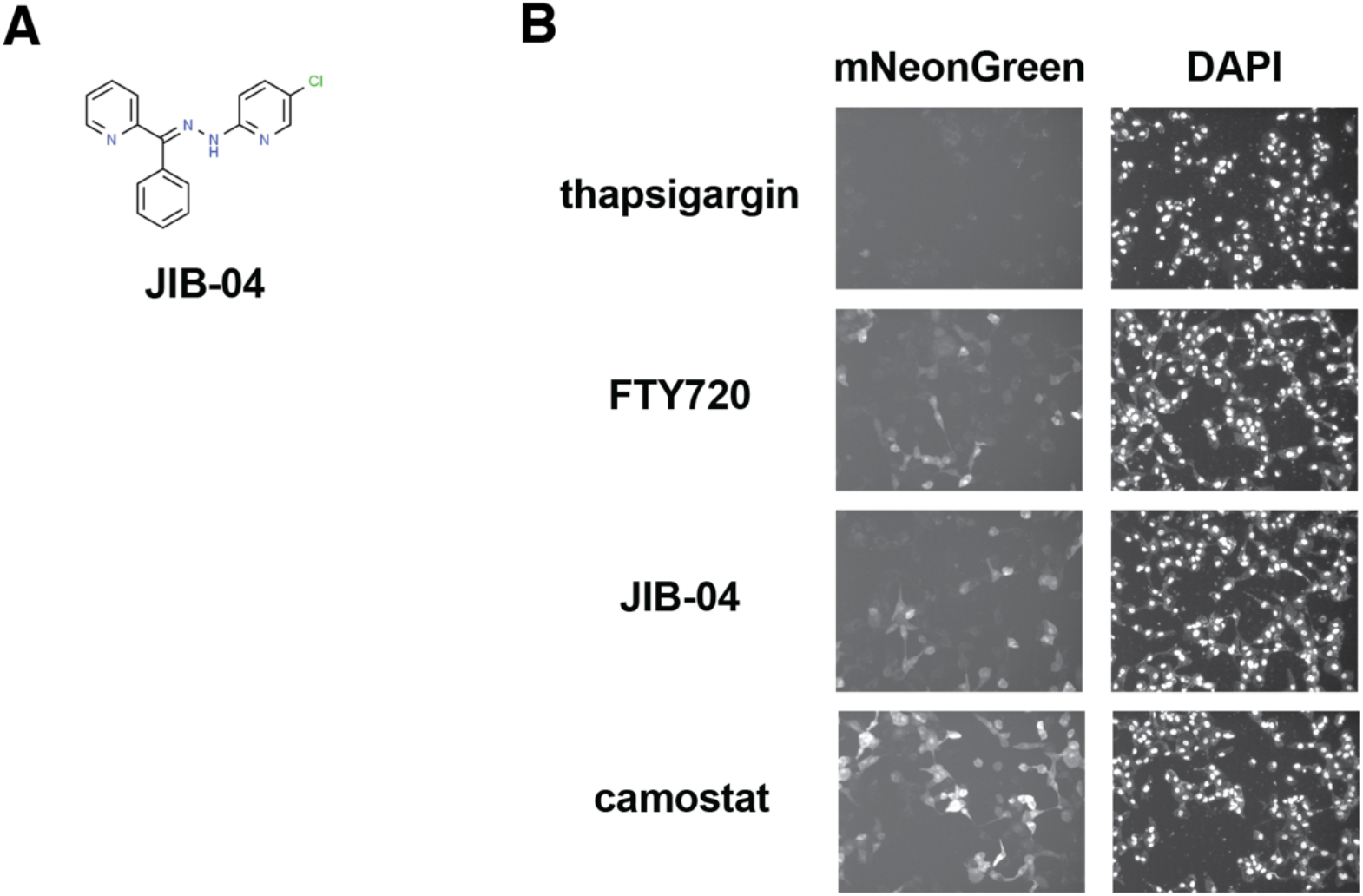
JIB-04 inhibits SARS-CoV-2 replication. (A) Chemical structures of JIB-04 E-isomer from ChemSpider database. (B) Representative images of Vero E6 cells infected by SARS-CoV-2-mNeonGreen (MOI=0.5) at 24 hpi in Fig. 1A.

**Fig. S2.**
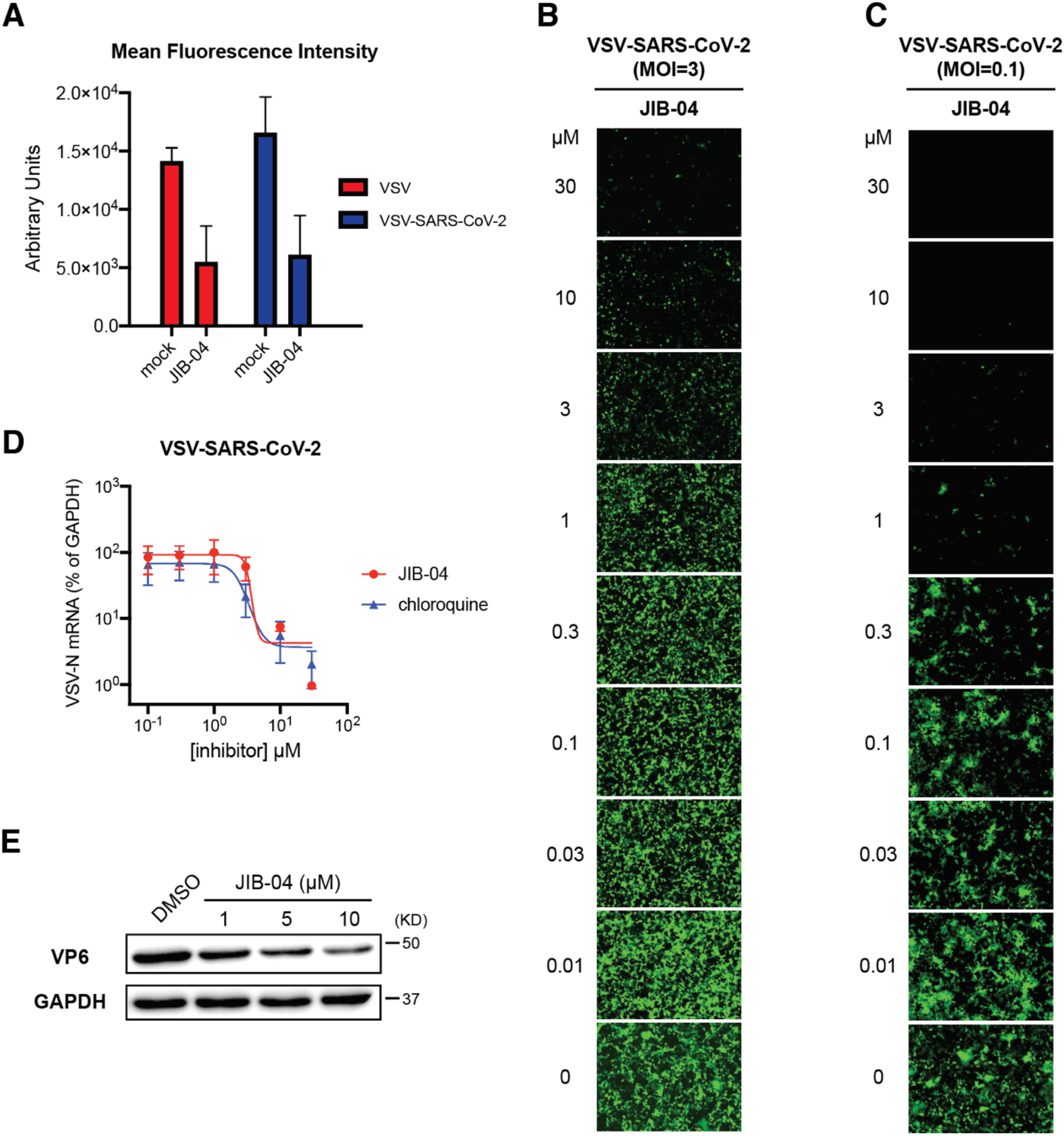
JIB-04 inhibits the replication of multiple viruses. (A) Mean fluorescence intensity of GFP positive cells in Fig. 2B was quantified by flow cytometry. (B) Dose-response analysis of VSV-SARS-CoV-2 replication with JIB-04 treatment. MA104 cells were treated with JIB-04 at indicated concentrations for 1 h and infected with VSV-SARS-CoV-2 (MOI=3). At 24 hpi, images of GFP positive infected cells were acquired by the ECHO fluorescence microscope. (C) Same as (B) except that cells were infected with an MOI of 0.1. (D) Dose-response analysis of intracellular viral RNA levels with JIB-04 or chloroquine treatment. MA104 cells were treated with compounds at 0.1 to 30 μM for 1 h and infected with VSV-SARS-CoV-2 (MOI=3). VSV RNA levels at 24 hpi were measured by RT-qPCR. (E) Western blot analysis of RV antigen VP6 levels with JIB-04 treatment. HEK293 cells were treated with JIB-04 at 1, 5, or 10 μM for 6 h and infected with porcine RV (MOI=0.01) for 12 h. GAPDH was used as a loading control. All experiments were repeated at least three times with similar results. Data are represented as mean ± SEM.

**Fig. S3.**
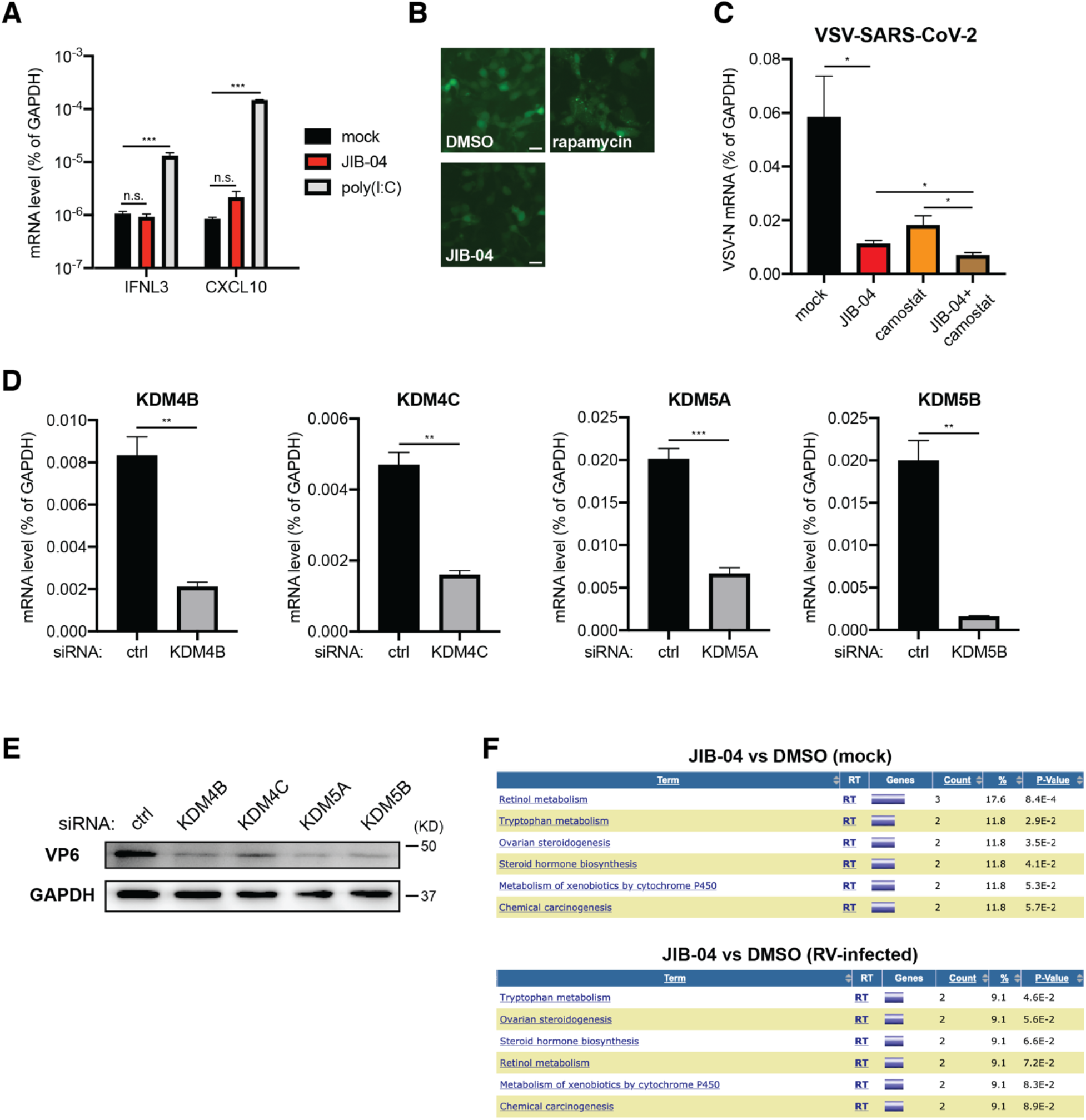
Inhibition or knockdown of specific KDM histone demethylases inhibits virus replication. (A) Expression of IFN and IFN-stimulated genes with JIB-04 treatment. HEK293 cells were treated with JIB-04 (3 μM) or transfected with low-molecular-weight poly(I:C) (100 ng/ml) for 24 h. mRNA levels of IFNL3 and CXCL10 were measured by RT-qPCR. (B) Autophagy formation with compound treatment. HEK293 cells were transfected with EGFP-LC3 plasmid for 24 h and treated with rapamycin (100 nM) or JIB-04 (3 μM) for another 18 h. GFP positive punctate structures indicate autophagy activation. Scale bar, 20 μm. (C) Intracellular viral RNA levels with JIB-04 and camostat treatment. Calu-3 cells were treated with compounds (10 μM) for 1 h and infected with VSV-SARS-CoV-2 (MOI=3). VSV RNA levels at 24 hpi were measured by RT-qPCR. (D) siRNA-mediated knockdown of JIB-04 target histone demethylases. HEK293 cells were transfected with scrambled siRNA or siRNA targeting indicated histone demethylases for 48 h. mRNA levels of indicated histone demethylases were measured by RT-qPCR. (E) Western blot analysis of RV antigen VP6 levels in cells with histone demethylase siRNA knockdown. HEK293 cells were transfected with scrambled siRNA or siRNA targeting indicated histone demethylases for 48 h and infected with porcine RV (MOI=0.01) for 12 h. (F) Pathway enrichment analysis of gene expression regulated by JIB-04 treatment. Downregulated genes in Fig. 3F with p values < 1e-10 were analyzed by DAVID functional annotation. For all panels except B, experiments were repeated at least three times with similar results. Fig. S2B was performed twice. Data are represented as mean ± SEM. Statistical significance is from pooled data of the multiple independent experiments (*p≤0.05; **p≤0.01; ***p≤0.001).

**Fig. S4.**
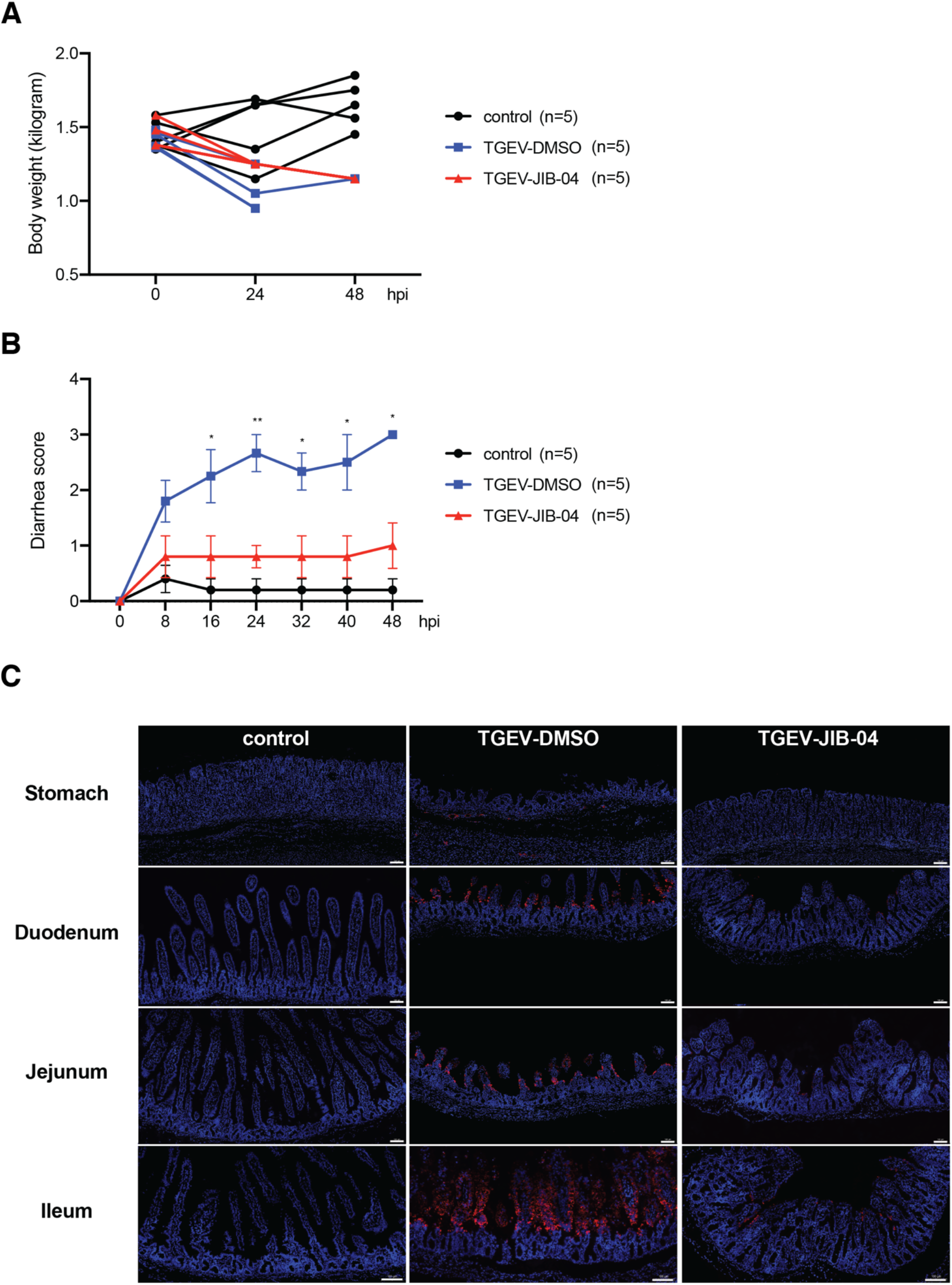
JIB-04 reduces TGEV induced weight loss and pathogenesis. (A) Weight of TGEV infected pigs with JIB-04 treatment in Fig. 4B. The body weight of individual animals was monitored every 24 h. (B) Diarrhea occurrence in TGEV infected pigs with JIB-04 treatment in Fig. 4B. Diarrhea severity was scored for the fecal specimens of DMSO or JIB-04 treated, mock or TGEV infected animals every 8 h. (C) Immunofluorescence staining of TGEV antigen in different GI tract sections from pigs sacrificed at 48 hpi. Blue: cell nuclei; red: TGEV nucleocapsid protein. Representative images of 3 animals. Scale bar, 100 μm.

## Author contributions

J.S., Q.Z., R.Z., and S.D. designed, executed, and analyzed *in vitro* efficacy studies. M.F.G.C., H.P.K., and G.H. assisted with the RNA extraction and RT-qPCR analysis. Y.Z. performed the *in vitro* TGEV inhibition studies. Z.L. performed the flow cytometry analysis. L.C. wrote the algorithm that quantifies inhibitor screen results. P.W.R. and S.P.J.W. constructed the VSV-SARS-CoV-2 virus. E.D.M. provided JIB-04 Z-isomer. Q.Z., J.Z., and R.G. propagated and titrated viruses. J.B.C. propagated and infected the clinical isolate of SARS-CoV-2. P.Y.S. provided the recombinant SARS-CoV-2 mNeonGreen virus. A.L.B propagated the mNeonGreen virus and designed the SARS-CoV-2 Taqman probe. S.H., B.L., and S.D. designed the *in vivo* efficacy studies. S.H., J.Z., X.C., B.F., and B.N., performed the *in vivo* TGEV infection experiments, dissected the animals and harvested tissues, and measured viral titers and cytokine mRNA levels. X.W., E.D.M., S.P.J.W., M.S.D., A.C.M.B., B.L., and S.D. provided supervision and funding for the study. J.S. and S.D. wrote the manuscript with the input and edits from S.H., Q.Z., J.B.C., J.Z., Z.L., M.F.G.C., H.P.K., G.H., S.P.J.W., M.S.D., A.C.M.B., and B.L.

## Acknowledgements

The authors thank Dr. Harry Greenberg (Stanford University, USA) and Dr. Bolívar A. Villacís-Bermeo (Guayaquil, Guayas, Ecuador) for constructive comments and arguments. We appreciate the assistance from Matthew Williams (Department of Molecular Microbiology Media and Glassware Facility) and Erica Lantelme (Department of Pathology and Immunology, Flow Cytometry Core Facility). Cytation plate scanning was assisted by Zhou Huang (Department of Molecular Microbiology).

## Data and materials availability

All raw data in the current study are available in Table S3 and Dataset S1. RNA-seq dataset has been deposited onto NCBI GEO (GSE156219).

